# A spatially localised cryoinjury model to study liver regeneration in zebrafish

**DOI:** 10.1101/2025.10.02.680167

**Authors:** Marcos Sande-Melon, Beth K. Logan, Andrew G. Cox, Juan Manuel González-Rosa

## Abstract

This Protocol Extension describes adapting an existing cryoinjury protocol to study heart regeneration in zebrafish^1^. We recently characterized a novel hepatic cryoinjury model that induces localized necrosis, inflammation, and fibrosis, which resolves as the liver regenerates. Here, we present a detailed description of the hepatic cryoinjury procedure. One of the key advantages of this model is that it allows direct comparison of healthy and injured liver parenchyma within individual animals. Our hepatic cryoinjury model provides a rapid and reproducible platform to investigate the molecular and cellular mechanisms underlying fibrosis and liver regeneration. This Protocol Extension aims to extend the reach of the cryoinjury approach for deployment in studying liver regeneration.

## Introduction

Unlike most organs, the human liver exhibits a remarkable ability to regenerate throughout life^2^. Despite this unique attribute, the molecular underpinnings that enable an injured liver to regenerate are still poorly understood^3^. Chronic liver diseases, including metabolic dysfunction-associated steatotic liver disease (MASLD) and fibrosis, are known to inhibit the regenerative capacity, leading to liver failure^4^. A better understanding of the mechanisms driving adaptive regeneration and maladaptive chronic liver disease will inform clinical scenarios associated with these complications and reduce the necessity of liver transplantation.

Liver regeneration is a multicellular response to an insult. Previous methods of liver injury, including partial hepatectomy (HPx)^5–7^, genetic ablation^8^, or hepatotoxins^9–24^, have been informative in identifying key regulators of liver repair in mice and zebrafish. We have recently developed the first spatially localised liver injury model in zebrafish, enabling the study of injured and healthy parenchyma^25^. Using this injury paradigm, based on cryoinjury (CI), we have characterised the development and regression of a fibrotic scar, the dynamics of liver cell proliferation, angiogenesis, and immune responses. Additionally, tissue-clearing approaches such as the advanced CUBIC method allow for deeper, three-dimensional imaging of the injured liver, providing novel insights into regenerative processes^26^. In contrast to other injury models, our hepatic CI model recapitulates critical features of liver pathophysiology observed in humans^25^. In the original protocol, a cardiac CI model that resembled myocardial infarction was established in zebrafish. Since publication, the cardiac CI model has become a gold standard for the study of heart regeneration^27–29^. We propose that the hepatic CI model is a cost-effective and rapid method to perform chemical and genetic screens to identify novel regulators of regeneration and fibrosis in adult zebrafish. Here, we describe a detailed protocol for hepatic CI in the zebrafish and provide guidance on liver dissection, collection, and preservation.

### Experimental design

Adult zebrafish are fasted from the evening before the surgery to reduce mortality associated with the procedure. On the day of the surgery, zebrafish are anesthetised by immersion in 0.032% (wt/vol) tricaine. Anesthetized fish are then positioned ventral side up on a sponge with an indentation, allowing for immobilisation and precise manipulations under a stereomicroscope. The cryoprobe is transferred to a liquid nitrogen container to cool the copper wire for at least two minutes. Ventral scales near the midline, between the pectoral fins, are gently removed using forceps, and the area is dried with tissue paper. To expose the liver while minimizing injury to adjacent organs, the skin is gently pulled using forceps, and a small midline incision is made using microdissection scissors. As a result, the ventral lobe of the zebrafish liver is exposed. To limit the cryoinjury (CI) to the liver, the exposed surface is gently dried with a cotton swab, avoiding the damaging blood vessels during this process. The cryoprobe is then quickly placed onto the surface of the liver until it thaws, approximately 15 seconds. Following the procedure, adult zebrafish are transferred to a tank and revived by flushing fresh water over the gills using a Pasteur pipette. The entire procedure should be completed in 2-5 minutes to ensure robust recovery from surgery **(Figure 1)**. The same procedure is performed on sham-operated animals, except the cryoprobe remains at room temperature. After surgery, animals are housed in standard zebrafish tanks with continuous flow. Operated animals may be fed the following day.

**Figure 1.**
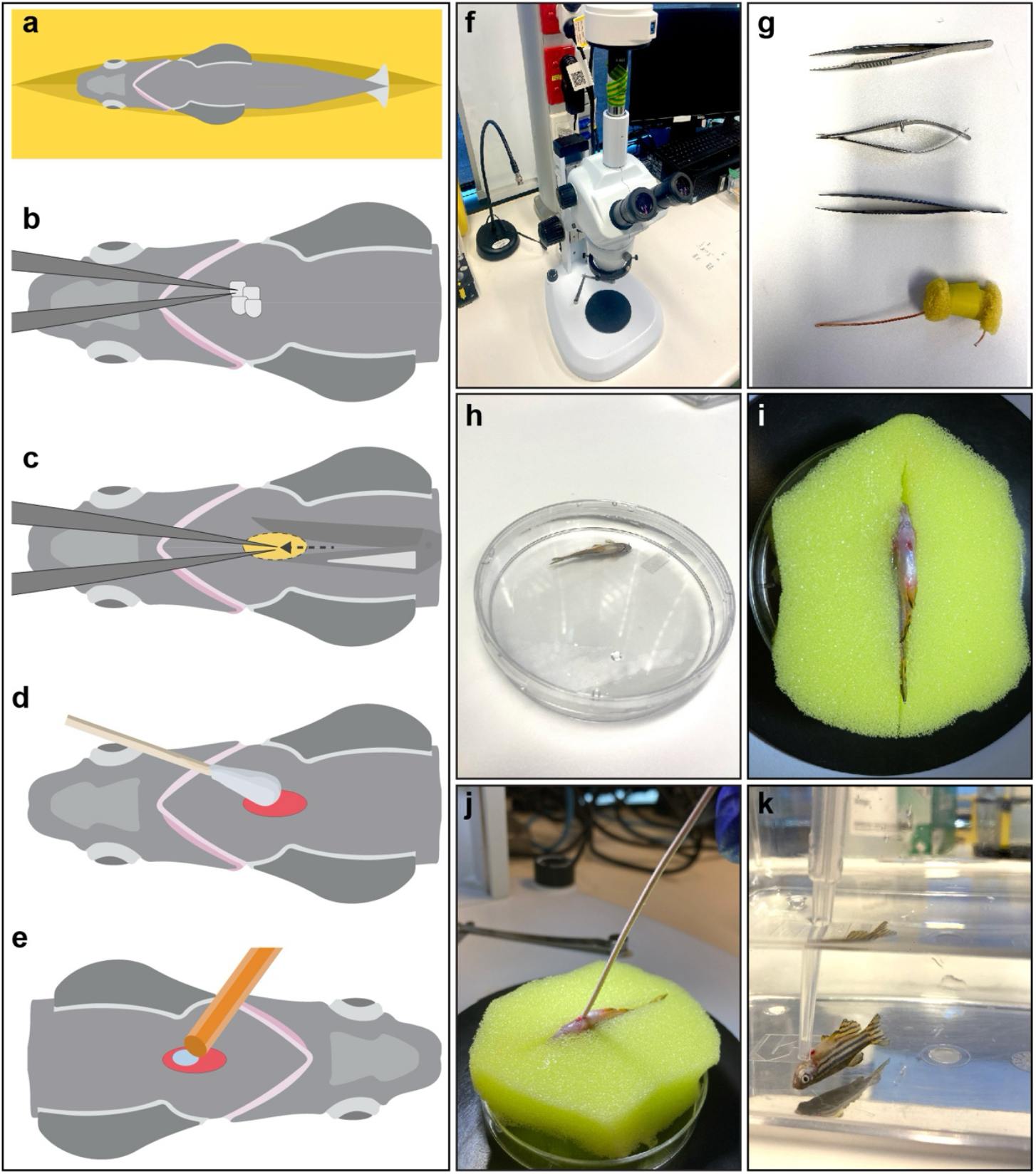
Steps to perform hepatic cryoinjury in the adult zebrafish. **(a)** Place the adult zebrafish in the sponge. **(b)** Remove scales from the area between the pectoral fins. **(c)** Perform a small incision in the ventral area to expose the liver using microdissection scissors, then pinch the skin to prevent internal damage to the liver (yellow highlighted area). **(d)** Dry the liver surface with a cotton swab. **(e)** Place the frozen cryoprobe on the liver surface. **(f)** A stereomicroscope is requiered to perform the hepatic cryoinjury. **(g)** Necessary surgical tools for zebrafish liver injury. **(h)** Place the adult zebrafish in tricaine water in a petri dish. **(i)** Adult zebrafish after ventral resection with the liver exposed. **(j)** Cryoinjury of the zebrafish liver. **(k)** Zebrafish reanimation using a plastic Pasteur pipette to increase the flow of fresh water throughout the gills in a recovery tank.

**Figure 2.**
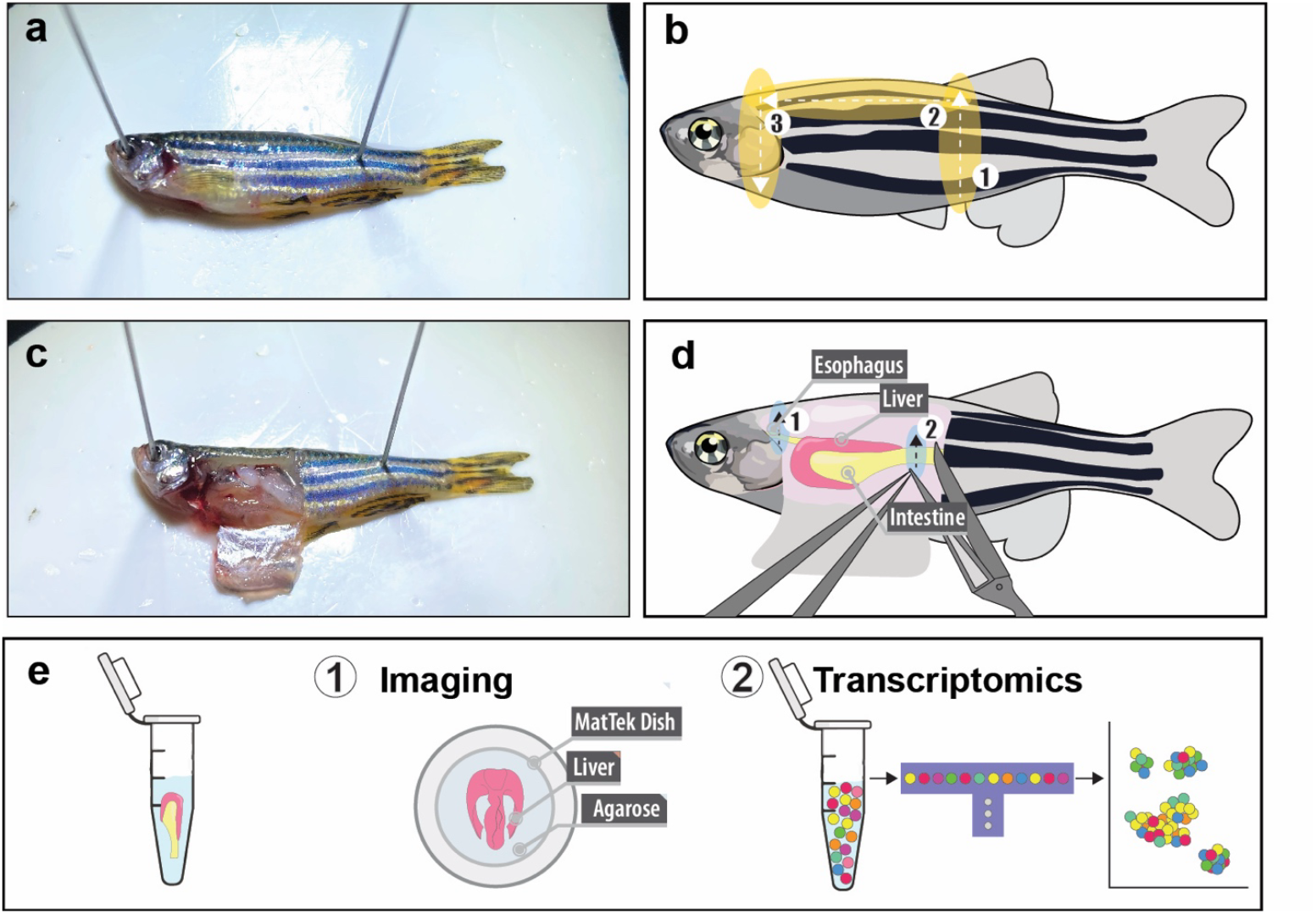
Liver collection after hepatic cryoinjury. **(a)** Mount the adult zebrafish on a silicone base and secure it with two pins. **(b)** Schematic of the incision used to expose the peritoneal cavity. **(c)** Once the peritoneal cavity is exposed, the gastrointestinal tract and liver can be easily visualized. **(d)** Schematic of the two necessary incisions to collect the liver and gastrointestinal tract. **(e)** Collected livers can be fixed and used for whole-mount imaging, or used for transcriptomics analysis.

**Figure 3.**
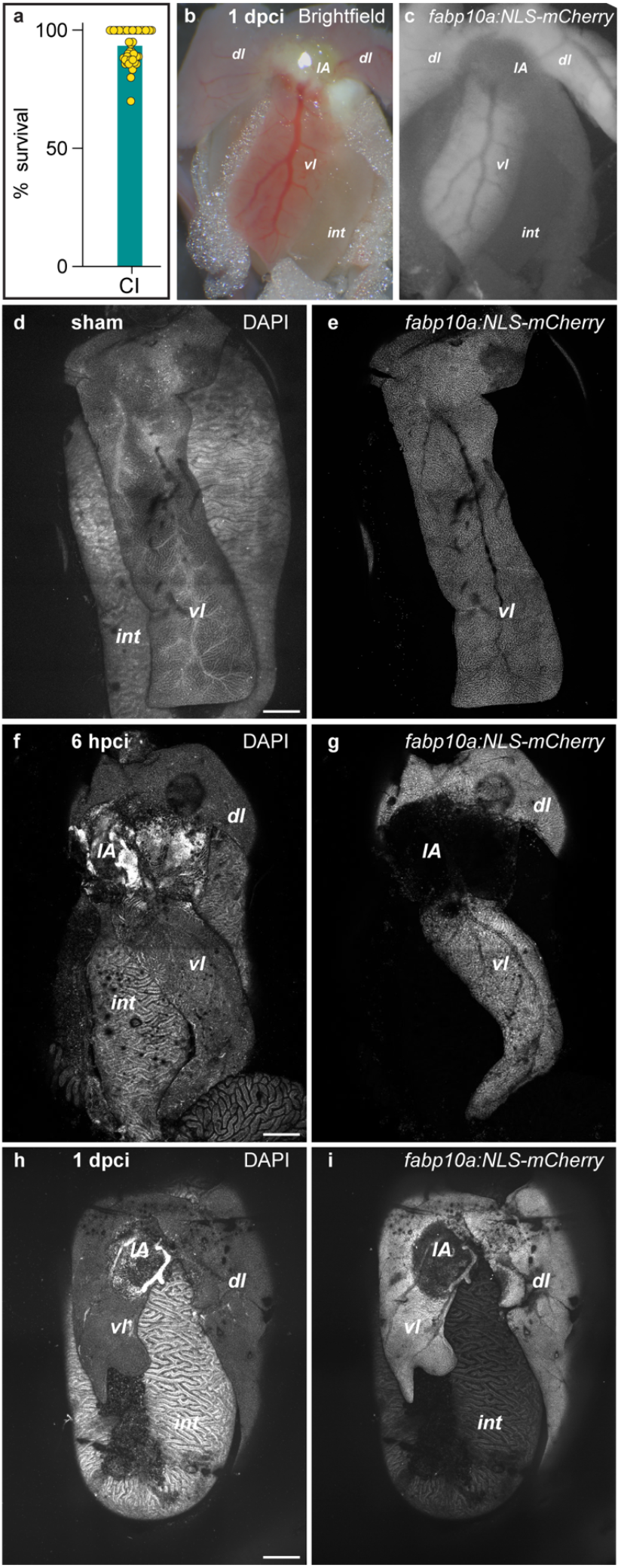
Imaging the extent of regeneration following hepatic cryoinjury in adult zebrafish. **(a)** Survival percentage after liver cryoinjury (n= 444; survival rate= 92.97%). **(b)** Brightfield image of an adult zebrafish liver at 1 day post-cryoinjury (dpci), showing the injured area. **(c)** In animals carrying an hepatocyte-specific fluorescence reporter, the cryoinjured area appears as an area that lacks mCherry^+^ signal. **(d-i)** Adult livers from sham-operated (d,e) or cryoinjured animals carrying the *Tg(fabp10a: NLS-mCherry)* transgene, at the indicated times after injury. dl: dorsal lobe; IA: injured area; int: intestine; vl: ventricular lobe.

**Figure 4.**
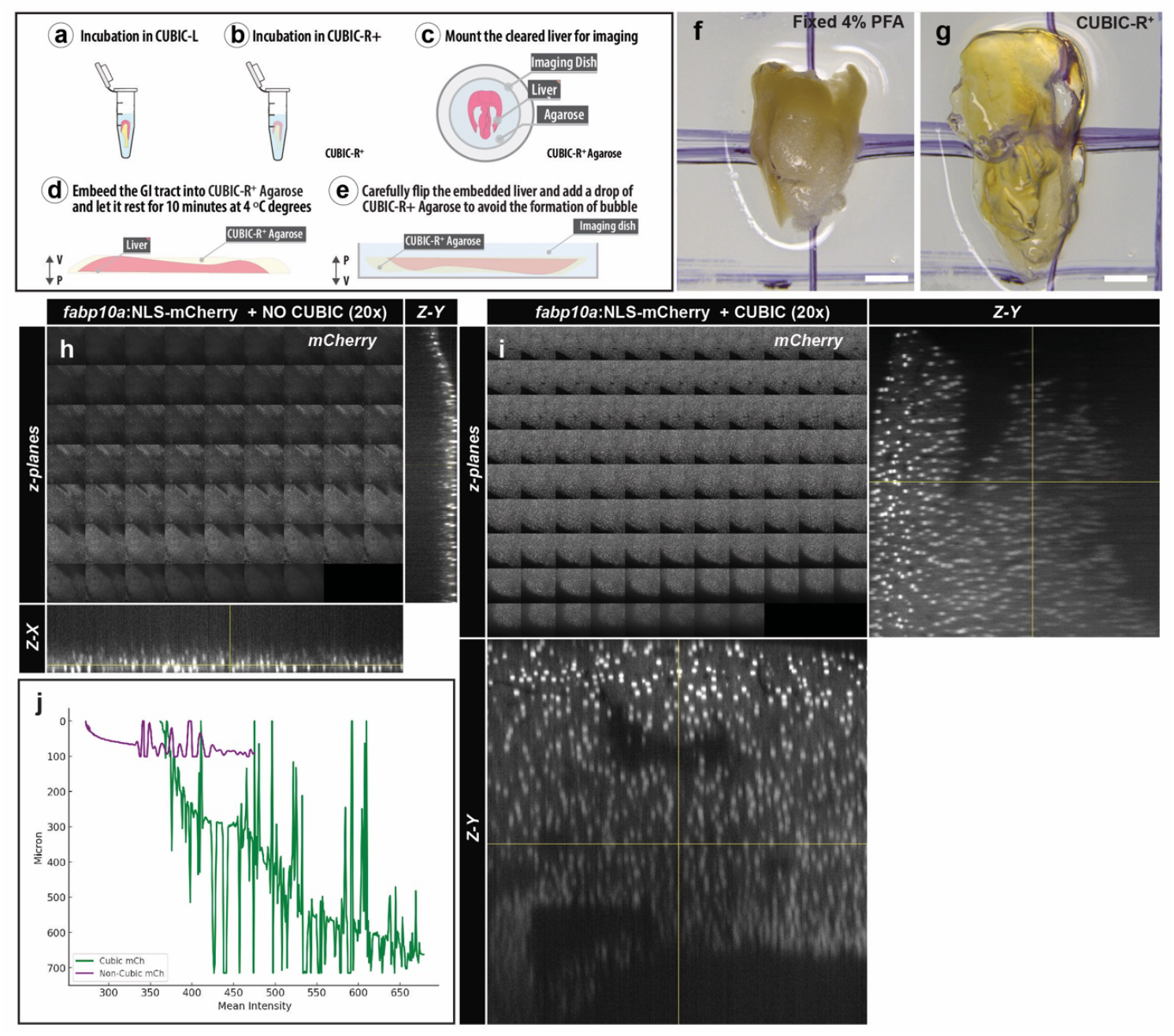
CUBIC-based clearing of the adult zebrafish liver enhances deep-tissue fluorescence imaging. **(a–e)** Diagram of the sample preparation protocol. **(a)** Incubate fixed livers in CUBIC-L, **(b)** transfer to CUBIC-R^+^, **(c)** mount the cleared liver in the imaging dish, **(d)** embed the gastrointestinal (GI) tract or liver in CUBIC-R^+^ agarose, and **(e)** carefully top off the agarose to prevent bubble formation before imaging. **(f, g)** Macroscopic view of an adult zebrafish liver before (f) and after (g) CUBIC-R^+^ clearing. The clearing process makes the tissue translucent. Scale bars = 1.1 mm. **(h, i)** Representative confocal z-stack images of adult zebrafish livers expressing the *fabp10a:NLS-mCherry* reporter without (h) or with (i) CUBIC clearing. Each panel shows successive optical sections (z-planes), with orthogonal cross-sections (Z–X and Z–Y) shown on the right and bottom, respectively. **(j)** Quantification of mean mCherry fluorescence intensity (x-axis) versus imaging depth (y-axis) for CUBIC-cleared (green) and non-cleared (purple) samples, demonstrating improved signal retention in deeper planes after clearing.

To assess liver regeneration, animals are euthanised and their livers harvested at the desired time point post-cryoinjury. Ethical euthanasia can be performed by immersing zebrafish in 0.16% tricaine. Livers can then be collected along with the entire gastrointestinal (GI) tract by making a transverse incision on the oesophagus and intestine. Livers should be harvested in ice-cold PBS for transcriptomic analysis or transferred to the desired fixative (10% Neutral Buffered Formalin, NBF; or 4% Paraformaldehyde, PFA) as soon as possible.

The extent of the injury in adult zebrafish can be observed as an opaque white blister using a bright-field stereomicroscope or as the lack of fluorescence when using hepatocyte-specific reporters such as *Tg(fabp10a:NLS-mCherry)*. Tissue damage can be detected using cell death detection methods such as TUNEL staining or anti-cleaved caspase-3. Alternatively, samples can be sectioned and subjected to histological staining. For example, we have successfully identified the injured area using the Acid Fuchsin Orange G (AFOG) and hematoxylin-eosin stainings^25^.

The hepatic cryoinjury regeneration model is highly reproducible, especially when comparing fish of the same gender and age. Notably, the zebrafish liver exhibits sexual dimorphism, with the female liver being larger than its male counterparts. Therefore, the injured area must be calculated as a percentage of the total liver parenchyma. The variety of liver-specific transgenic animals, combined with new high-throughput techniques, enables the timely study of specific liver populations using this method. Given the economic advantages and easy manipulation of the zebrafish, we expect the hepatic cryoinjury model to be a rapid and reproducible pipeline for drug discovery in regenerative medicine.

### Advantages and limitations of hepatic cryoinjury in zebrafish

The hepatic cryoinjury (CI) model is the first to enable precise, spatially localised injury, allowing the study of the microenvironmental differences between the healthy and injured liver parenchyma within the same animal. As a result, this model allows the study of the interactions between the healthy and injured parenchyma at the injury border during inflammation, fibrosis, and regeneration. Importantly, hepatic cryoinjury recapitulates key pathophysiology features associated with human chronic liver diseases. Unlike other liver injury paradigms such as HPx, hepatic CI induces massive cell death and fibrosis, which resolve as the liver regenerates. Cell-specific ablation has also been used to induce liver injury, and this method has enhanced our understanding of the role of specific cell types during liver regeneration. However, these models required complex transgenic backgrounds that are lineage-restricted and therefore less capable of accurately modelling liver disease. In contrast, the hepatic CI model induces cell death in hepatocytes, biliary epithelial cells, and endothelial cells, resembling focal liver necrosis in humans. Overall, our protocol for hepatic CI in zebrafish has the potential to deepen our understanding of the molecular mechanisms behind regeneration and fibrosis. This knowledge is essential to understanding liver disease and identifying potential regenerative strategies.

The age, size of the animals and the relatively moderate throughput of the method are the main limitations of the CI model. We have successfully induced liver injuries in animals ranging from 3 to 12 months post-fertilization. Although juvenile injuries may be possible by adjusting the size of the cryoprobe, this method is not applicable to studying regeneration in embryonic stages. Moreover, the procedure’s efficiency and reproducibility depend on surgical expertise and are thus subjected to a learning curve.

In contrast to the embryonic samples, analyzing adult samples may be particularly challenging. To solve this limitation, we have integrated in this protocol efficient tissue-clearing approaches, such as advanced CUBIC, which enables high- resolution imaging of the injury site and surrounding liver architecture. Through CUBIC-based clearing, otherwise opaque regions of the damaged liver become optically transparent, allowing deep-tissue visualization of fluorescent reporters and subcellular structures. These improvements enable more precise quantification of injury progression, cellular dynamics, and regenerative events during critical post-injury stages. However, some limitations should be considered. First, the CUBIC clearing process is time and reagent-intensive, requiring multiple incubation steps that extend the experimental timeline. Second, although fluorescent signals are generally well preserved in CUBIC-cleared samples, some spectral shifts or uneven bleaching can occur when working with specific fluorescent proteins or dyes. Third, the application of CUBIC clearing is currently limited to fixed tissue preparations and thus does not permit live imaging of injury or regeneration in live zebrafish. Finally, as with any tissue-clearing method, specialized imaging equipment, including long-working distance objectives, is necessary to fully exploit the optical transparency of the organ. Nonetheless, when optimized, CUBIC clearing significantly enhances ability to study hepatic regeneration in the zebrafish liver, providing anatomically intact, 3D views of cellular processes critical for understanding tissue recovery.

## Procedure

### Hepatic Cryoinjury

1. Ensure the zebrafish are fasted starting the evening before surgery.
2. Place the zebrafish tank at the surgery table and allow the animals to settle for at least 15 minutes before the surgery.
3. Prepare a new tank with zebrafish system water to reanimate the adult zebrafish after surgery. Place the tank into an incubator at 28 °C until the surgery commences.
4. Prepare the anaesthetic mixture by diluting tricaine to a 0.032% concentration in zebrafish water using a 50 mL Falcon tube and a plastic Pasteur pipette. Pour the anaesthetic medium into a new Petri dish.
5. Clean the microdissection scissors and sharp forceps using 70% ethanol.
6. Wet the surgery sponge with anaesthetic medium and squeeze out the excess water. Place the wet sponge in a petri dish lid, focus the stereomicroscope, and adjust the illumination to improve visibility during surgery.
7. Use a clean tissue paper or cotton swabs to dry the liver surface during the procedure.
8. Pre-cool the cryoprobe for at least 2 minutes in liquid nitrogen. Dry the cryoprobe between surgeries and time accordingly to ensure consistent results.
9. Immerse the adult zebrafish in the anaesthetic medium.
10. Once the fish is anesthetized, which usually happens within a minute, gently pinch the caudal tail to check for reflexes. If there is no response, carefully grasp the tail with forceps and place the fish on the damp sponge prepared for surgery. Position the animal ventral side up, with its head facing left.
11. Use the stereomicroscope to focus and locate the pectoral fins. Descale the medial area between the fins.
12. Make a longitudinal incision in the ventral region between the pectoral fins to expose the ventral lobe of the liver.
13. Carefully dry the liver surface with a cotton swab or tissue. If the animals carry a tissue-specific reporter, you can use a fluorescence stereomicroscope to better localise the liver.
14. Quickly place the cooled cryoprobe on the liver surface for 15 seconds. After this, the injured area will appear as an opaque white blister.
15. Use forceps to gently grasp the fish by the tail and return it to the recovery tank.
16. Flow water over the gills with a plastic Pasteur pipette to facilitate rapid recovery from anaesthesia and improve survival rates. During recovery, ensure fish are kept at 5 fish per liter to maintain appropriate density.
17. Place the zebrafish in their corresponding racks in the fish facility and maintain them until harvesting.

### Liver Harvesting

18. Euthanise adult zebrafish by immersing them in 0.16% tricaine diluted in zebrafish water.
19. Place the adult zebrafish on the dissection bed inside a Petri dish lid, with the ventral side facing up and the head positioned to the left, as previously described. Use pins to secure both edges, ensuring stability for collecting the gastrointestinal (GI) tract.
20. Open the peritoneal cavity by making three incisions. Firstly, cut vertically from the perianal fins towards the dorsal area. Next, make a second longitudinal incision from the end of the first incision towards the head, cutting through the thoracic cavity. The third incision starts from the endpoint of the second incision and continues towards the pectoral fins. This creates a peritoneal window that exposes the liver and gastrointestinal (GI) tract.
21. Using microdissection scissors, cut the oesophagus and the intestine without damaging the liver.
22. Collect the liver together with the GI using forceps.
23. Fix the liver with the GI tract intact for histology in 4% PFA at 4 °C. Avoid over-fixing the samples, as this can increase autofluorescence. Adult zebrafish livers should not be immersed in 4% PFA for more than 24 hours to ensure optimal results.
24. Alternatively, the freshly isolated liver can be snap-frozen in an Eppendorf tube immersed in liquid nitrogen for further genomic or transcriptomic analysis.
25. Wash the livers three times for 20 minutes in 1x PBS-Tween20 upon fixation. If not processed immediately, immerse the samples in 30% sucrose diluted in PBS 1X overnight at 4 °C, placing them vertically in a Falcon tube. Transfer the livers to a plastic mold filled with OCT and place it on dry ice. Once the OCT is completely frozen, store the samples at –80 °C in Ziplock bags until further use.

### Liver Clearing

26. If the livers are going to be cleared, transfer them to CUBIC-L for 18 h at 37 °C. Use a hybridisation oven with rocking agitation.
27. After CUBIC-L incubation, wash the livers in 1x PBS-Tween20 three times for 15 min.
28. Incubate the livers in blocking solution (5% BSA, 5% goat serum, 0.1% Tween-20) at 4 °C with rocking agitation.
29. Primary and secondary antibodies should be incubated for 24 hours each at 4 °C with rocking agitation.
30. Transfer the livers into CUBIC-R+ for 2 h at room temperature. Protect the samples from the light to prevent photobleaching of the fluorophores.
31. Embed the livers in freshly prepared 1% CUBIC-R+-Agarose for imaging.
32. Transfer the livers onto a glass-bottom culture dish (MatTek Corporation) for confocal acquisition. Orient the sample for imaging.
33. After the agarose has solidified, flip the sample and add a drop of CUBIC-R+-Agarose. This step ensures that the injured area is positioned closest to the objective, ensuring it will be the first region imaged in an inverted microscopes.
34. Image the livers on the desired microscope.

## Materials

- Zebrafish. Adult zebrafish males from 4 to 9 months old. Experiments were approved by the Peter MacCallum Cancer Centre AEEC (E665) and the Institutional Animal Care and Use Committees of the Massachusetts General Hospital and Boston College. All animal procedures conformed in accordance with the Prevention of Cruelty to Animals Act, 1986 (the Act), associated Regulations (2019) and the Australian code for the care and use of animals for scientific purposes, 8th edition, 2013 (the Code).
- Zebrafish water (animal zebrafish facility aquatic system).
- Copper of 1 mm thickness (Jaycar; Australia).
- Liquid nitrogen. Wear adequate PPE and handle carefully.
- Liquid nitrogen container.
- Cryoprobe
- Paraformaldehyde (PFA, 16% (wt/vol); Electron Microscopy Sciences, cat. no. 15710.
- Phosphate-buffered saline (PBS 1x).
- Tricaine (Ethyl 3-aminobenzoate methanesulfonate salt, Merck no. A-5040).
- Plastic Pasteur pipette (3 ml; Deltalab, cat. no. 200037).
- 50 mL Plastic tube (50 ml; Falcon, cat. no. 352098).
- Absorbent cellulose tissues (KimWipe).
- Dissecting stereomicroscope (Leica M165FC fluorescent stereomicroscope or similar).
- Microdissection scissors (Spring scissors, 8-mm blades; Fine Science Tools).
- Forceps (Dumont no. 5; Fine Science Tools).
- Petri dish (BD Falcon, cat. no. 353003).
- Agarose (Sigma-Aldrich, cat. no. A9539).
- N-Butyldiethanolamine (Sigma-Aldrich, cat. no. 471240).
- 2,3-dimethyl-1-phenyl-5-pyrazolone (antipyrine; Sigma-Aldrich, cat. no. A5882).
- Nicotinamide (Sigma-Aldrich, cat. no. 72345)
- N,N,Nʹ,Nʹ-Tetrakis(2-hydroxypropyl)ethylenediamine (Sigma-Aldrich, cat. no. 122262)
- Urea (Sigma-Aldrich, cat. no. U5378)
- Imidazole (Sigma-Aldrich, cat. no. I5513)
- Triton X-100 (Sigma-Aldrich, cat. no. T8787).

### Reagent set up

- PFA 4% (wt/vol). Dilute 10 ml of 16% PFA in 40 ml of PBS 1X. Aliquot and store at − 20 °C until use.
- Tricaine 0.4% (wt/vol). Dissolve 2 g of Tricaine in 500 mL of fish water. Mix it by vigorously shaking it in a 1L bottle. The bottle must be covered with aluminium foil to avoid Tricaine degradation.
- Sponge. Perform a partial incision in the sponge sufficient to hold the adult zebrafish steadily.
- Cryoprobe. 5 cm of 1 mm thickness Copper of 1mm diameter was cut. The copper was passed through a sponge and fixed with electric tape.
- CUBIC-L: Mix 10% (wt/wt) N-butyldiethanolamine and 10% (wt/wt) Triton X-100 in distilled H2O with a stirrer at room temperature (22–26 °C). CUBIC-L can be stored at room temperature for up to 6 months.
- CUBIC-R+: Mix 45% (wt/wt) antipyrine, 30% (wt/wt) nicotinamide, 0.5% (vol/vol) N-butyldiethanolamine and distilled H2O. CUBIC-R+ can be stored at room temperature for up to 6 months.
- 1% CUBIC-R+ Agarose: Mix 2% (wt/wt) agarose powder with CUBIC-R+ at room temperature, ensuring the solution is well mixed through vigorous stirring. Heat the mixture in a microwave until boiling starts. Carefully remove the bottle, securely cap it, and shake well. Repeat the heating and shaking steps multiple times until the agarose is fully dissolved. 1% CUBIC-R+ Agarose can be stored at room temperature for up to 6 months.

## Troubleshooting

### Surgery timing

Steps 1-6: 15 minutes.

Steps 7-16: 2-5 minutes.

Steps 17-22: 10 minutes.

Step 25: 60 minutes.

### High Mortality

- Consider the age and health of the adult zebrafish used for surgery. Animals between 4 and 9 months generally show higher survival rates and yild a more consistent injury area.
- The anaesthesia mixture should be prepared with precision, carefully pipetting the correct tricaine concentration into the appropriate volume of zebrafish water. Before surgery, test the anesthetic solution by briefly immersing a healthy, uninjured zebrafish and verify full recovery. If recovery is delayed or incomplete, reassess the water quality, prepare a fresh anaesthesia mixture, and replace the water in the recovery tank.
- Regularly check the water quality parameters in the zebrafish racks. Fluctuations in temperature, nitrate levels, or pH can significantly impact zebrafish health and reduce survival during surgical procedures.
- Replace the cryoprobe after every 100 uses to maintain its reliability and optimal performance.

### CUBIC Liver Clearing

- Samples embedded in CUBIC-Agarose and stored at 4 °C for several hours may form crystals. To avoid crystallization, remove the samples from agarose and transfer them into a suitable storage medium. If crystal appear, heat the solution to 60 °C until they fully dissolve.
- Prolonged incubation in CUBIC-L at 37°C can cause tissue degradation. Carefully monitor and limit incubation time.
- Extended exposure to CUBIC-R+ can decrease sample fluorescence intensity; minimize exposure duration to preserve signal.

### Reproducibility

- Use a timer to ensure precise and consistent cryoprobe chilling between surgeries.
- Position the probe vertically to maintain a uniform contact surface across all procedures.
- Ensure that zebrafish are maintained at consistent densities in the facility tanks.
- Utilise zebrafish of the same age and sex to ensure consistency and standardize experimental conditions.
- Liver sexual dimorphism is pronounced between male and female; account for this variable when designing experiments.
- Feed the animals on the morning following surgery, ensuring that feeding schedules remain consistent across all experimental procedures and harvesting times.

### Anticipated results

Hepatic CI has a survival rate of ∼93% (n= 444)^25^. Liver injury can be observed at 1 day post-cryoinjury (dpci) in the parenchyma as a distinct white necrotic area accompanied by loss of fluorescence when using hepatocyte zebrafish reporters. Upon hepatic CI, a necrotic core surrounded by an annular zone of apoptosis can be detected at the site of injury. Loss of hepatocytes (HCs) and biliary epithelial cells (BECs), the two main cell types of the liver parenchyma, is evident at 1 dpci. Endothelial cells are also adversely affected by hepatic CI. Hepatocyte and BEC proliferation peaks between 3 and 7 dpci, whereas the vasculature is largely restored by 7 dpci.

A transient scar is formed during the first 7 dpci and progressively regresses until its fully resolved by 30 dpci. A strong immunological response occurs during the first 7 dpci, returning to baseline levels by 30 dpci. Implementation of a CUBIC-based tissue clearing protocol can further enhance the visualization of these necrotic, fibrotic, and regenerative processes in 3D, providing deeper insights into the cellular and structural dynamics of liver repair. We anticipate that the hepatic CI model will be a powerful approach to study liver cell death, fibrosis formation and resolution, hepatocyte and biliary epithelial proliferation, inflammatory responses, and vascular regeneration.

## Acknowledgments

M.S.-M. is supported by the Early.Postdoc Mobility fellowship from the Swiss National Science Foundation (SNSF). J.M.G.-R. is supported by the National Institutes of Health (1R01HL164749-01), the American Heart Association (19CDA34660207), the Corrigan Minehan Foundation (SPARK Award), and the Hassenfeld Foundation (Hassenfeld Research Scholar Award). A.G.C is supported by an NHMRC Investigator Grant (GNT1176650) and an ARC Discovery Project Grant (DP200102693). We also extend our thanks to the Peter MacCallum Cancer Centre Core Facilities and their staff who supported this work, namely the Centre for Advanced Histology & Microscopy, the Molecular Genomics Core, Flow Cytometry and Bioinformatics Core Facilities. We also thank the staff involved at the University of Melbourne Zebrafish Core Facility (DRUM) and Biology Optical Microscopy Platform (BOMP). Finally, we thank members of the Cox Laboratory (Peter MacCallum Cancer Centre), the González-Rosa Laboratory and the Organogenesis and Cancer Program at the Peter MacCallum Cancer Centre for the helpful discussions.

## Author contributions

M.S-M and B.L performed experiments. M.S-M, J.M.G.-R. and A.G.C designed experiments and prepared the manuscript.

